# Investigating the existence of periodicity in activity of neural network by novel neural signal processing technique - quantifying induced learning in cell culture

**DOI:** 10.1101/177360

**Authors:** Sayan Biswas

## Abstract

The network forming ability of neurons are huge for their sparking ability to form new connections and break existing ones. This sheer ability allows dynamic nature of the network for which this network are ever changing. The neurons being cells that are chemically and electrically excitable, electrical excitation of these cells cause variation of voltage in vicinity of the active neurons. These variation captured through electrical recording device records to activity points in the network. Cultured neuron cells on Multi electrode array dish is used to study disassociated cultures. A novel integrative model of neural signal processing termed as Activity Index is applied. AI variation is plotted graphically to show the evidence in periodicity of network analysis. The finding on periodicity are discussed along with how could it be used as a potential parameter to quantify learning ability of a cell culture.

**Index Terms:** neurons, dynamic, variation of voltage, Multi electrode array, Activity Index train, Periodicity, learning

## I. INTRODUCTION

The first studies about brain dates back to the scientist Cajal [1]. Since then researchers are curious in finding how actually neurons are physically connected in nervous system and what is the significance of such a connection in functional neuronal network [2] [3] [4]. It has been seen these functional connectivity in nervous system are reasons to cognitive and behavioural states in brain. Brain potentially has high parallel processing [5] [6] abilities against the silicon chips. Brain receives many inputs from surrounding but essentially attends [7] [8] selectively to the important essential inputs. This selection of inputs from many possible one requires huge computations by brain. Such computations [9] are carried out by brain in network of neurons. Neurons forms network [10] by joining [11] each other through their synapses.

Cultured dissociated neurons on the micro electrode array-MEA [12] [13] is a very powerful method for understanding the functional and structural characteristics of in-vitro biological neuronal networks. This powerful device has helped researchers to investigate the network level mechanisms occurring in the cultured neurons. Such investigation has played a crucial role in understanding the cellular basis of learning [14], memory [15] [16] and plasticity [17] in the synapses. MEA has evidently allowed researchers to carry out recording extending over months which has certain advantages like easier control and less complexity than comparable in vivo systems. Research involving MEA can be helpful in designing of hybrid systems [18] or understanding brain models. Hybrid systems is a result of combination of artificial and natural intelligence. MEA technology can be used to investigate the interaction occurring between live neuron and artificial systems to build hybrid systems. Such hybrid systems could be used to stimulate and study different pathological situations like epilepsy or stroke. MEA can contribute in understanding brain models by allowing researchers to understand structural and functional connection in brain contributing in natural intelligence of brain. MEA is implemented in real time experiments [19] where a feed back loop is used by delivering electrical stimulus. These are also called closed loop experiments. These electrical stimulation to the neuron network helps to control network activity and also induce synaptic plasticity to it.

## II. MATERIAL AND METHODS

The data for computational analysis is obtained from the public repository [20]. The cortical cell of the culture is obtained from embryonic wisar rats. The cortex is kept in appropriate temperature and pressure in the required living medium and solution for keeping the cells alive. These cultures of cells are made to keep in the MEA dish from which recording of electrical activity is obtained. Neurons are electrically and chemically excitable cells. The neurons during its active state fires a action potential [21] which exists for 5ms to 10ms. During this firing process the neuron has a variation of voltage occurring across it membrane. These variation of voltage occurs only when the neuron is in active state. These active state remains for a short period of 5ms to 10 ms. As the variation of voltage occurs across the membrane of the neuron there happens to be all total variation of voltage in the vicinity of the active neurons. This variation of voltage or potential occurring in the vicinity of active electrodes is called Extracellular field potentials-EFP [22] [23].

These EFP is captured through the electrodes of the MEA dish. A generic recording paradigm is illustrated in figure 1. On high pass filtering this EFP neural spike train is obtained. Neural spike train (figure 3) is recorded for all the 60 recording electrodes. Neural spike train gives the precise time of occurrences of spikes in the neural network. These time points of activity at each electrode was available through the available data set. One important point to be noted is all electrodes of the MEA don’t participate in recording activity. Some are grounded with respect to which the potential or EFP is recorded on the other electrodes.

**Fig. 1:**
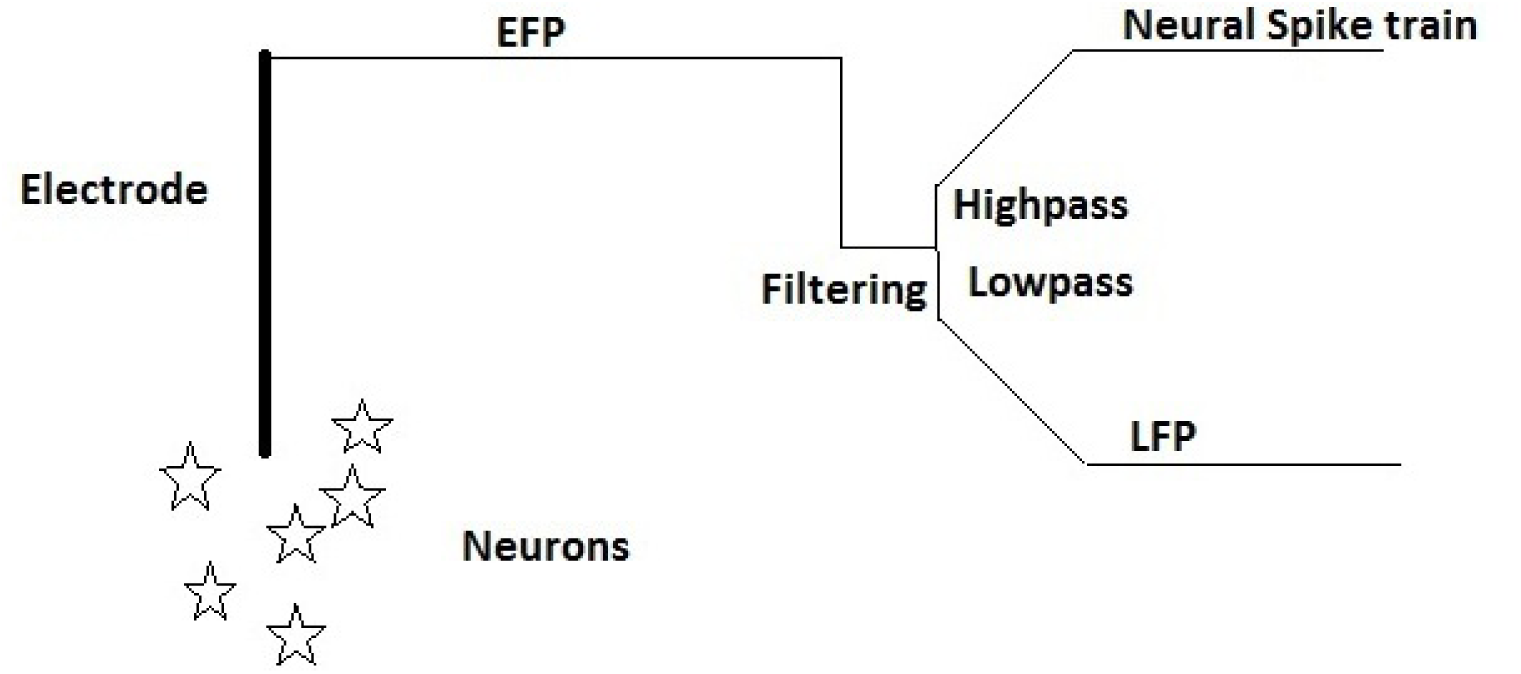
A generic recording paradigm.

**Fig. 3:**
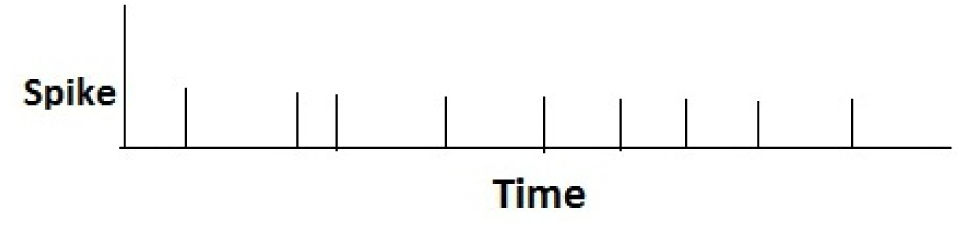
Neural spike train.

## III. ACTIVITY INDEX-NEURAL SIGNAL PROCESSING

METHOD

Activity index - AI is a parameter obtained from analysis of neural spike train data analysis. This parameter essentially plays a critical role in analysis of a dynamic neural network. If *S*(*a, t*_1_*, t*_2_) denotes the number of spikes fired f rom e lectrode a i n t he t ime w indow [*t*_1_*, t*_2_], AI of electrode a at time point t is calculated as

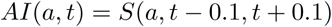

Activity index of all the electrode at two random time point s illustrated in figure 4. A ctivity i ndex values a ttained by a particular electrode at all time points is called Activity index train (figure 2). C omputing A I i s t he n ovel neural signal processing method implemented in this manuscript. Importance of the parameter AI is discussed below:

- To understand dynamic network activity.
- Aimed to signify network status by accounting effect of traveling spike.
- Comments about activity transfer(or electric signal transfer) in the network.

**Fig. 2:**
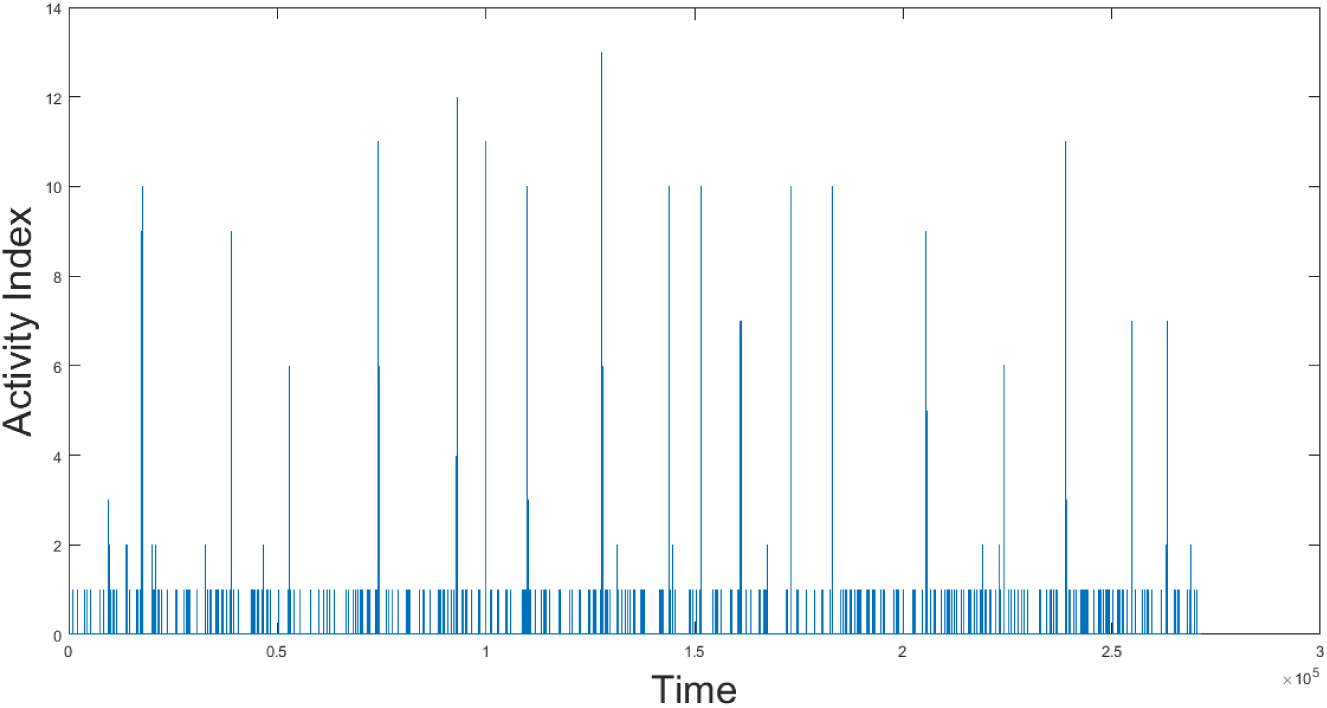
Activity Index train.

**Fig. 4:**
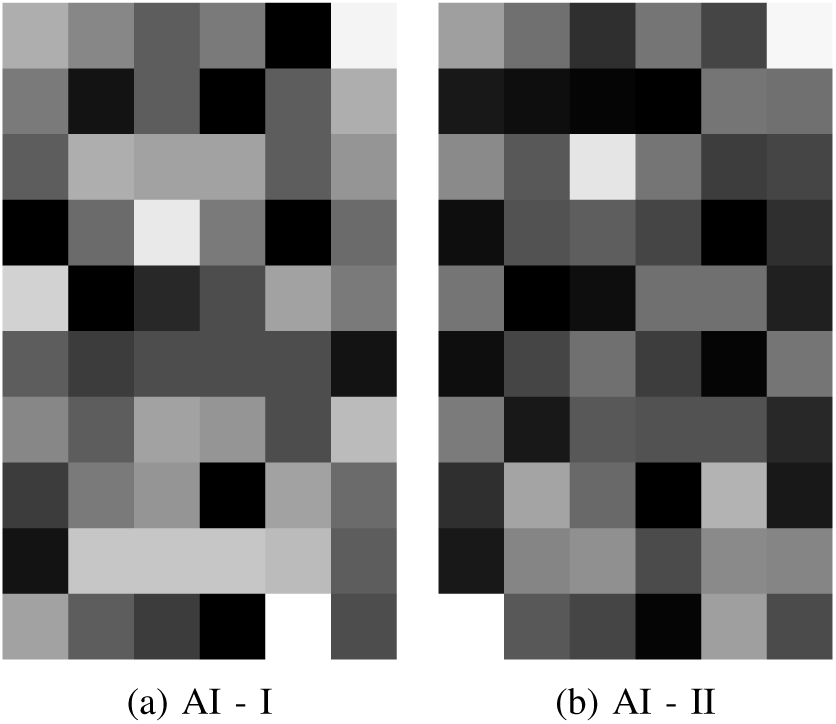
Activity Index.

Earlier work on Activity index dealt with ranking of nodes [24] and commented regarding the information available [25] [26] from neural nodes. Earlier work has a detailed treatment of Activity Index and Activity Index train explaining the computation of this new neural signal processing method.

## IV. PERIODICTY

Periodic function are those that repeat values after a certain intervals. The repetition might occur in spatial domain, temporal domain or spatio temporal domain. A function *f* is periodic with period *T* if,

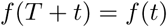

. The function *f* will repeat its values after definite interval of time T or more importantly fundamental period *T*. Essentially periodicity property of a signal can be determined by looking by a visual analysis of a signal. If a specific contour is repeated after specific periods then it must be periodic.

## V. PERIODICITY IN THE NEURONAL NETWORK

From the data set of neural spike train AI train for each of the electrode was computed. The Activity Index AI for each electrode is plotted against the time. Some of these plots are shown in figure 5. In all these plots data of 20 electrode over 100000 time point is shown. On visual analysis it could be found that AI rise was occurring in a network during particular time frames, when all electrode began to have non zero AI values. Such occurrences of all electrode’s rising AI values at particular time can be termed as time of rising activity. During this time the liveliness in the network reaches high. This happens after certainly not periodic intervals but after intervals which can be assumed to be imperfectly periodic or almost periodic. Looking at the shape of the variation of AI over electrode at the time of rising activity it could be noted that the contours are of similar shapes. This idea can be safely approximated as the AI value of a electrode during rise time are approximately same which results in similar contour of AI variation at the rise time. To summarise the finding:

- Each electrodes has a characteristic AI value. During rise time electrodes has AI value near to its characteristic value.
- Similar contour of AI variation during time of rising activity.

**Fig. 5:**
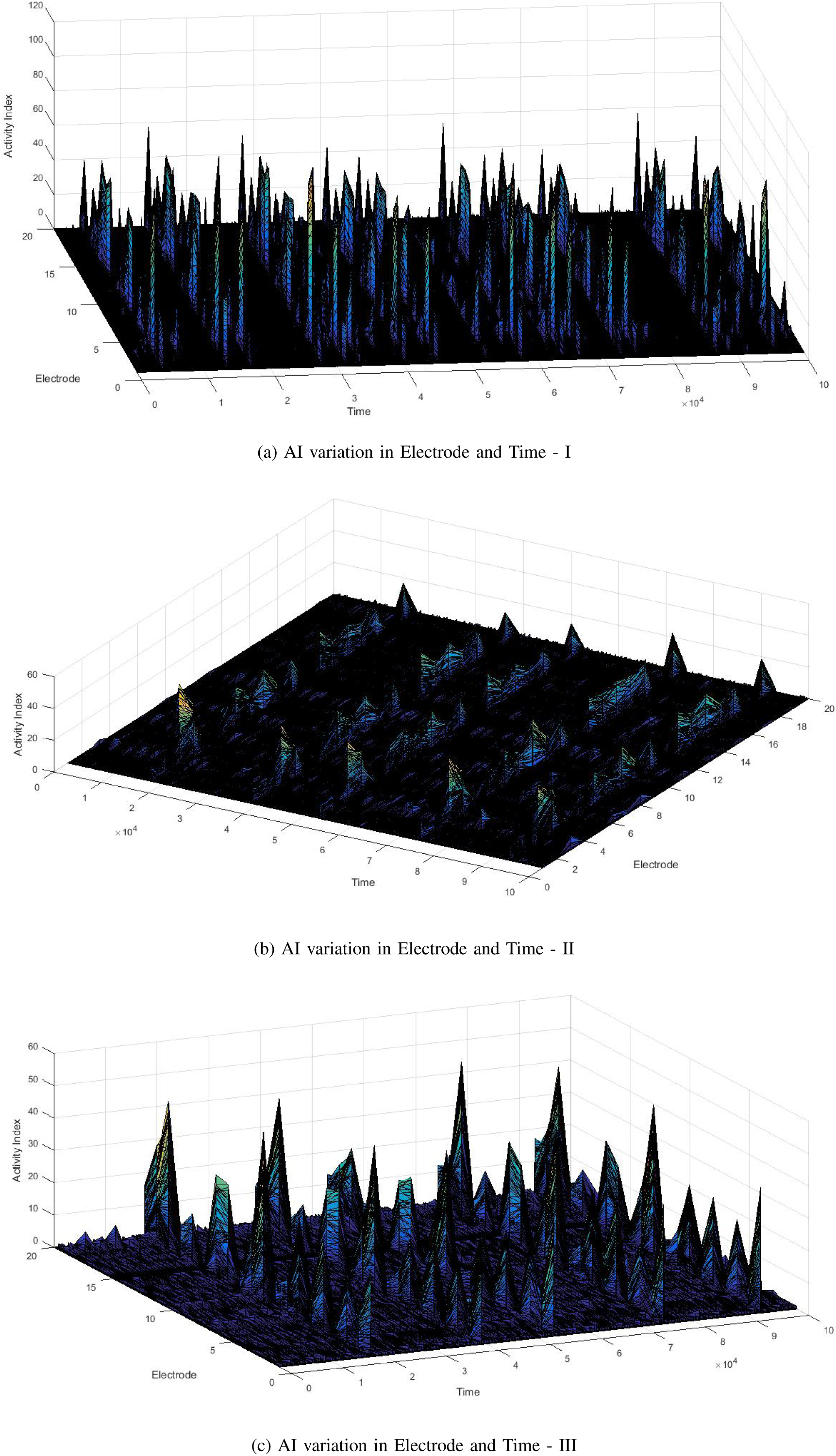
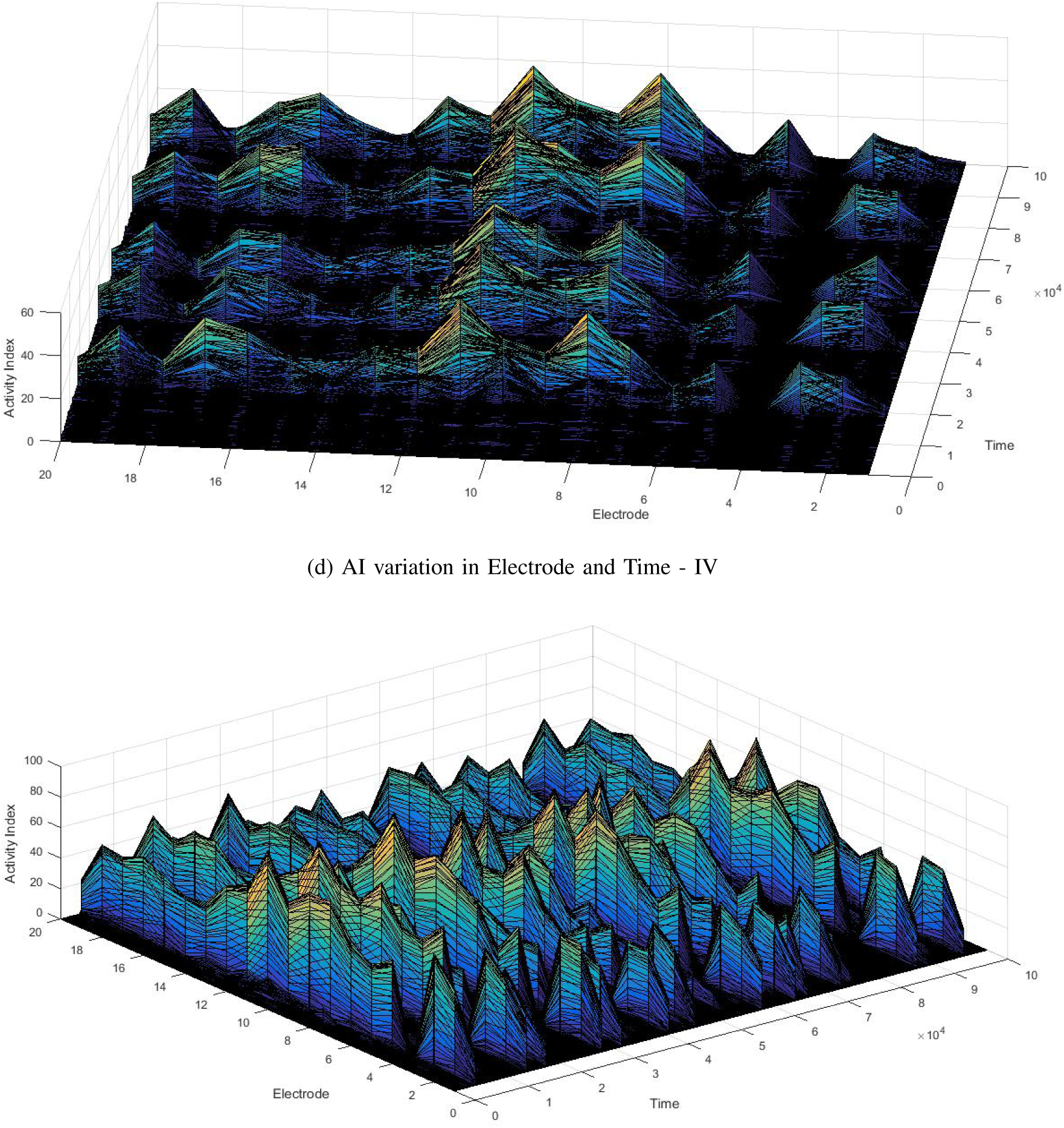
AI variation in Electrode and time of 5 different trial.

## VI. INDUCED LEARNING

Brain in which neuronal network process information is able to learn information from external sources. This external source can be thought to be as the learning media. This learning occurs in the neuron network itself. Induced learning in the in-vitro neuron culture is possible by the stimulating certain electrodes of the MEA dish by voltage. Added to this information how could the learning in the network be parameterised is helpful. The timing difference between consecutive time of rising activity might prove to be a parameter of learning. How well a parameter could learn could be effectively understood by the extent by which the difference between time of rising activity could be modulated. Hence difference between time of rising activity could be a parameter for understanding learning ability of a network.

## VII. DISCUSSION AND CONCLUSION

This work finds that the variation of AI in time and electrodes can be considered roughly periodic. As AI-Activity Index proves as a parameter to understand the dynamic network activity, it was evidenced that in the network activity rises and falls together. As an application of the signals being roughly periodic, network activity roughly repeats after specific intervals. Hence the need for using an entire recording for purpose of analysis could be reduced to analysing the signal for specific time period, which could roughly be two or three periods of the signal. Hence the finding of periodicity could be of great application when it comes to analysing a network based on its Activity Index. Such a finding could greatly reduce the computational burden. Hence by analysing data of two three periods, overall network activity could be satisfactorily understood.

It could also be investigated on how can the periodicity, or the intervals between two consecutive time of rising activity in the network, serve as a evidence of quantity of learning possible through network. As a part of this investigation different part of the network could be stimulated with a pulsating voltage and hence induce synaptic plasticity in the network forming stronger connection in between part of network. Following to which it could be seen how can such variation in the network plasticity cause the periodicity to vary hence quantifying how well network is able to learn.

As stated it was seen from the visual plot that the AI value with respect to a electrode is almost kind of similar for every rise times. Hence at every rise time an electrode took a AI value not very different from its value at the earlier rise time. This was elucidated earlier by calling every electrode has a characteristic value. To the knowledge of the author this happens as the first work where a quantity (AI) is found which characterises a electrode in the dynamic network.

## VIII. ACKNOWLEDGEMENT

Author would like to thank Jadavpur University, Department of Electrical Engineering.

